# Developing a standardized but extendable framework to increase the findability of infectious disease datasets

**DOI:** 10.1101/2022.10.10.511492

**Authors:** Ginger Tsueng, Marco A. Alvarado Cano, José Bento, Candice Czech, Mengjia Kang, Lars Pache, Luke V. Rasmussen, Tor C. Savidge, Justin Starren, Qinglong Wu, Jiwen Xin, Michael R. Yeaman, Xinghua Zhou, Andrew I. Su, Chunlei Wu, Liliana Brown, Reed S. Shabman, Laura D. Hughes, the NIAID Systems Biology Data Dissemination Working Group

## Abstract

Biomedical datasets are increasing in size, stored in many repositories, and face challenges in FAIRness (findability, accessibility, interoperability, reusability). As a Consortium of infectious disease researchers from 15 Centers, we aim to adopt open science practices to promote transparency, encourage reproducibility, and accelerate research advances through data reuse. To improve FAIRness of our datasets and computational tools, we evaluated metadata standards across established biomedical data repositories. The vast majority do not adhere to a single standard, such as Schema.org, which is widely-adopted by generalist repositories. Consequently, datasets in these repositories are not findable in aggregation projects like Google Dataset Search. We alleviated this gap by creating a reusable metadata schema based on Schema.org and catalogued nearly 400 datasets and computational tools we collected. The approach is easily reusable to create schemas interoperable with community standards, but customized to a particular context. Our approach enabled data discovery, increased the reusability of datasets from a large research consortium, and accelerated research. Lastly, we discuss ongoing challenges with FAIRness beyond discoverability.

## Introduction

Code and data sharing is becoming the norm in the biomedical sciences as scientists promote open science to facilitate reproducibility, to enable collaborative science and meta-analyses, and to maximize research investments from sponsoring agencies. As a result, an increasing number of publishers^1–6^ and funders^7–11^ require data management plans, open source code, and open access to data upon publication. Many of these requirements are codified in the improved NIH Data Sharing Policy, which goes into effect 25 January 2023^7,8^. To fulfill these mandates, a sea of biomedical research repositories have sprung up, creating a data sharing landscape that is difficult and time-consuming to navigate. In many cases, compliance is met by simply stating that data would be available upon request even without any mechanisms for enforcing data availability^12–14^. In order for research data and code to be shared and reused effectively, they must be Findable, Accessible, Interoperable, and Reusable (FAIR)^15^. However, findability requires useful, complete descriptions of the contents, known as metadata. Metadata allows researchers to discover and evaluate whether the dataset or code is suited for their purposes, provides helpful context for third party analysis, and enables sponsoring agencies to track compliance with open data policies. Comprehensive metadata are critical to facilitate the understanding, use and management of data and to support reproducible research^16–19^.

A lack of standardization of metadata is a primary hurdle in making research data and code more adherent to FAIR practice. Numerous biomedical researchers and domain-specific consortia have tried to address the standardization issue, resulting in a proliferation of best practices and data sharing recommendations^19–31^, which have contributed to confusion on which standards to adopt and how to implement their adoption. Even when standards are adopted, the standardized structured metadata is often unexposed and not reusable. The proliferation and fragmentation of incomplete data repositories, lack of organization of data in endless Data Lakes^32^ or in repositories with insufficient metadata, and the lack of common metadata standards make it difficult to combine separate data resources into a single searchable index. While standardizing metadata will not be sufficient to fully combine research data and code from different sources and enable meta-analyses, it is nevertheless a crucial first step towards this goal.

Encouraging metadata standardization has improved the findability of information on the web and enabled more informative displays and intuitive visualization of search results. Schema.org is an open-source project supported by Google, Microsoft, Yahoo, and Yandex to develop vocabularies to describe information across the internet. By creating a communal set of standards for information identification and organization, web crawlers can automatically access data and use it for a specific purpose. For instance, Google Dataset Search attempts to create an easy-to-search interface to find data by indexing datasets across the web that expose Schema.org-compliant metadata through HTML markup^33–35^. Analysis of such datasets can enable discoveries undetected in an individual dataset, including genotypic, epigenetic, phenotypic and epidemiologic correlates of disease or immunity to disease to improve human health. To this end, the NIH supports efforts to enhance biomedical data FAIRness and encourages the use of open domain-specific repositories, generalist repositories, and knowledge bases for broad sharing of data with the larger research community^22^. While some NIH-recommended repositories use Schema.org standards, Google Dataset Search does not yet fulfill the needs of biomedical researchers for reasons that include: the limitations of a broad schema not developed for biological data, divergence in formatting, coding and units of experimental data, limited features in the user interface, and lack of an Application Programming Interface (API). Ongoing efforts such as Bioschemas ^36,37^ exist to create more biologically-relevant schemas, but those efforts currently focus on developing properties across many different researchers and working groups to be integrated within Schema.org which may need to be further customized to a particular research context. Given these issues, the specific metadata standardization needs of National Institute of Allergy and Infectious Diseases (NIAID)- sponsored researchers, and the increasing number of available research datasets, we developed a Dataset and software (ComputationalTool) schema tailored for infectious and immune-mediated disease research.

Here, we describe our approach to develop this schema to aid in cataloging datasets across diverse research areas with shared interests in infection and immunity. Our goal was to develop a reusable process and framework that is compatible with large-scale FAIR data efforts, tailored to infectious and immunological disease datasets, and that is easy-to-adopt and extend to lower the barrier for metadata collection and secondary analysis. First, we performed a landscape analysis to understand how pervasive Schema.org standards are within the biomedical community and compare how these schemas are used. Next, we applied what we learned to develop a Dataset and ComputationalTool schema tailored for research outputs in the infectious disease research space. To help researchers easily collect and publish these metadata, we developed a guide to help identify dataset repositories and extended a form-based mechanism to easily collect dataset metadata based on our proposed standards when no community repository exists. In developing the minimalistic schema for infectious and immunological disease research, we demonstrate a process for improving domainspecific data FAIRness that may be generalizable. Our process is broadly extendable to other domains of biomedical research and serves as a reusable process for other research communities to create interoperable schemas that help make their research outputs more FAIR.

## Results

### Fragmented application of schemas results in inconsistent use of standardized properties

First, we sought to understand how widespread Schema.org standards are in common data repositories. We found that general purpose repositories, sources not limited to biological data, were more likely than specialized repositories to use Schema.org-compliant markup (**Figure 1a**). The broader data sharing community, represented by 78% of generalist sources surveyed, has coalesced on the Schema.org standard, but biological repositories lag in migrating to a single data standard. Although some more specialized sites included Schema.org markup, they were in the minority, with only 18% of repositories surveyed using the Schema.org standard. As a result, the vast majority of biomedical data repositories are neither findable in dataset projects such as Google Dataset Search, nor interoperable with dataset metadata from other sources.

**Figure 1.**
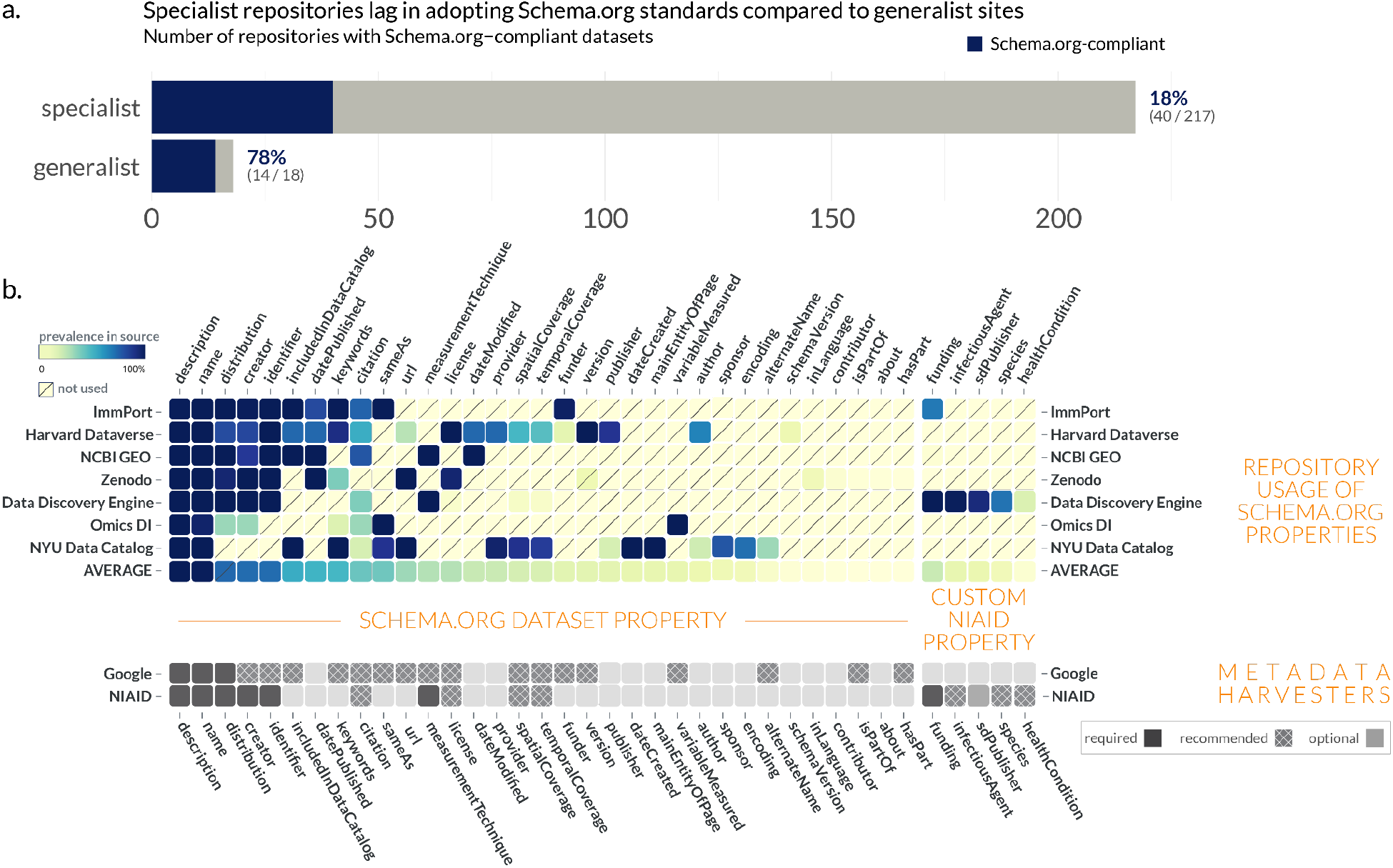
Distribution of Schema.org standards in common data repositories. **a**, Schema.org compliance in common biomedical repositories (see <u>Supplemental Table 1</u>). **b**, Within Schema.org-compliant repositories, each source used the standard differently. While description and name were almost universally provided for each dataset, other metadata properties were more inconsistently used within and between sources. Google Dataset Search marginality is provided in Ref. ^38^

To further investigate how Schema.org Dataset profile was being applied by general-purpose and specialized repositories, we compared the property utilization/prevalence across eight sources of biomedical datasets (**Figure 1b**). While Schema.org does define a set of properties to describe datasets, it is less specific regarding the structure (data type) of these properties or their relative importance. Indeed, the Schema.org Dataset specifies 124 properties, but in our survey of 7 different repositories, only 34 (27.4%) of the properties were used at all.

The adoption of the Schema.org Dataset properties was associated with usage by metadata harvesters. Name and description were almost universally provided, and are the only two fields required by Google Dataset Search. Of the remaining available properties, very few are actually relevant to promoting findability in biomedical datasets. In fact, the Bioschemas community selected only 13 (10.4%) out of 124 available properties from Schema.org’s Dataset as useful for describing a Dataset in the life sciences^39^. This may simply be a reflection of the fact that the Schema.org Dataset profile derives and inherits properties from the Schema.org CreativeWork profile, which has many general properties irrelevant to biological sciences. The prevalence of values for a particular property illustrates how these repositories prioritize certain properties over others. For example, ImmPort and Omics DI prioritize the sameAs property, which tracks either datasets cross-listed on other sites or reference web pages, compared to Zenodo, or Harvard Dataverse. Even if a property is shared between two different repositories, they may implement the property differently. For instance, is a publisher a string (“`NIAID Data Dissemination Working Group`”)or an object (`{name: “NIAID Data Dissemination Working Group”, sponsor: “NIAID”}`)? Is it a single publisher or many? Moreover, what is meant by `publisher` – the dataset collector(s), the host lab, the host institution, or the repository which publishes the data? This diversity – both in structure and semantic meaning – complicates efforts to find datasets between sources. Thus, analyzing the prevalence is helpful to inform the creation of our own schema by highlighting commonly used properties and resolving ambiguities for selecting between two semantically-equivalent properties (*e.g*. publication vs. citation).

### Designing a Dataset schema based on our analysis of Schema.org usage

Based on our analysis of commonly used Schema.org Dataset properties (**Figure 1b**), we developed a minimalistic-but-extendable schema tailored for infectious disease datasets. The NIAID Division of Microbiology and Infectious Diseases (DMID) Systems Biology (***NIAID SysBio***) Dataset schema contains 7 required fields, 7 recommended fields, and 1 optional field (**Table 1**). The schema uses Schema.org as a base and introduces four main changes: (***1***) the addition of infectious disease-specific properties, encompassing those pertaining to the pathogen and/or the host; (***2***) the addition of whether a property is required, recommended, or optional (*marginality*); and (***3***) a more defined structure for the properties, including more specific types, whether one or many values are allowed (*cardinality)*, and (***4***) linking to ontologies and controlled vocabularies to define what values are allowed for that property. By extending the Schema.org standard, we focus the metadata on properties tailored for immune-mediated and infectious diseases, while also allowing these datasets to be findable in Schema.org-based data discovery projects like Google Dataset Search.

**Table 1.** Comparison of three common Dataset schema specifications: Schema.org, Google Datasets, and the infectious disease-specific NIAID SysBio Dataset. Properties required in the schema are denoted by red text with (*). Note that Schema.org has additional properties not included in either Google Datasets or NIAID SysBio schemas which are omitted for simplicity. Supplemental Table 2 provides an extended crosswalk between the NIAID SysBio Dataset schema and other commonly used schemas.

| property | Schema.org | Google | NIAID SysBio |
| --- | --- | --- | --- |
| alternateName | Text | Text | NA |
| citation | CreativeWork, Text | CreativeWork, Text | ScholarlyArticle |
| ↳ citation.name | Text | NA | Text |
| ↳ citation.url | URL | NA | Text* |
| creator | Organization, Person | Organization, Person | Organization, Person* |
| ↳ creator.name | Text | NA | Text* |
| description | Text | Text* | Text* |
| distribution | DataDownload | DataDownload | DataDownload* |
| ↳ distribution.contentUrl | URL | URL* | Text* |
| ↳ distribution.dateModified | Date, DateTime | NA | Text* |
| ↳ distribution.encodingFormat | Text, URL | Text, URL | NA |
| funder | NA | Organization, Person | NA |
| funding | Grant (*pending*) | NA | MonetaryGrant* |
| ↳ funding.description | Text | NA | Text |
| ↳ funding.funder | Organization, Person | NA | Organization* |
| ↳ funding.funder.name | Text | NA | Text* |
| ↳ funding.funder.parentOrganization | Organization | NA | Text |
| ↳ funding.identifier | Text | NA | Text* |
| ↳ funding.url | URL | NA | Text |
| hasPart | CreativeWork | Dataset, URL | NA |
| healthCondition | NA | NA | Text |
| identifier | PropertyValue, Text, URL | PropertyValue, Text, URL | Text* |
| includedInDataCatalog | DataCatalog | DataCatalog | NA |
| infectiousAgent | NA | NA | Text |
| isAccessibleForFree | NA | Boolean | NA |
| isPartOf | CreativeWork, URL | Dataset, URL | NA |
| keywords | Text, URL | Text | NA |
| license | CreativeWork, URL | CreativeWork, URL | Text |
| measurementTechnique | Text, URL | Text, URL | Text* |
| name | Text | Text* | Text* |
| sameAs | URL | URL | NA |
| sdPublisher | Organization, Person (*pending*) | NA | Organization, Person |
| spatialCoverage | Place | Place, Text | AdministrativeArea |
| ↳ spatialCoverage.administrativeType | NA | NA | Text |
| ↳ spatialCoverage.alternateName | Text | NA | Text |
| ↳ spatialCoverage.identifier | PropertyValue, Text, URL | NA | Text |
| ↳ spatialCoverage.locationType | NA | NA | Text |
| ↳ spatialCoverage.name | Text | NA | Text |
| species | NA | NA | Text |
| temporalCoverage | DateTime, Text, URL | Text | TemporalInterval |
| ↳ temporalCoverage.duration | NA | NA | Text |
| temporalCoverage.endDate | CreativeWorkSeason,<br>CreativeWorkSeries,<br>DatedMoneySpecification,<br>EducationalOccupationalProgram, Event,<br>MerchantReturnPolicySeasonalOverride, Role, Schedule | NA | Text |
| temporalCoverage.name | NA | NA | Text |
| temporalCoverage.startDate | CreativeWorkSeason,<br>CreativeWorkSeries,<br>DatedMoneySpecification,<br>EducationalOccupationalProgram, Event,<br>MerchantReturnPolicySeasonalOverride, Role, Schedule | NA | Text |
| temporalCoverage.temporalType | NA | NA | Text |
| url | URL | URL | NA |
| variableMeasured | PropertyValue, Text | PropertyValue, Text | NA |
| version | Number, Text | Number, Text | NA |

We prioritized including properties which were commonly used within the community already (**Figure 1b**). For instance, author and creator are two similar fields used to express authorship of a dataset; however, the community, including Google Dataset Search, tends to prefer creator. Additionally, based on our goal of reducing collection burden on the researcher to promote uptake, we focused on developing a parsimonious schema to capture a key set of defining properties for biological datasets without being exhaustive. Two aspirational properties which generally have been neglected, but are important in the context of infectious disease research were also included: `temporalCoverage` and spatialCoverage. We created four new properties that do not exist within Schema.org Dataset to customize the schema for our needs. We added one required property, funding (now a pending property in Schema.org), and three optional infectious disease specific properties: `infectiousAgent, healthCondition`, and species. To encourage the capture of metadata provenance, we included the pending property `sdPublisher`, which tracks the original generator of metadata. Since funding is an essential property to allow funders to track research outputs and for data users to understand provenance, we define the source of funding as a required property. While available in Schema.org, funding information is not generally captured by repositories, and some providers like Harvard Dataverse collect funding information^40^ but do not standardize or expose it^41^. To promote compatibility with other commonly used schemas, we provide a crosswalk between our NIAID SysBio schema, Schema.org, and Google Dataset Search (**Table 1**), as well as a more extensive table (**Supplemental Table 2**) which compares our schema to an analysis of data schemas including DataCite and Dublin Core by the RDA Research Metadata Schemas Working Group^42^.

### Developing a ComputationalTool schema to link datasets and software

To derive optimal meaning from datasets and promote reanalysis, we wanted to be able to facilitate linkage of datasets to the computational tools and software used to generate and analyze them. In order to increase the findability of computational tools, we reviewed Schema.org’s SoftwareSourceCode schema and the Bioschemas ComputationalTool profile to generate a minimal set of properties to describe and link relevant computational tools. This schema was extended from the Bioschemas ComputationalTool and has 6 required, 10 recommended, and 4 optional properties (**Table 2**). As with the Dataset schema, the ComputationalTool schema includes additional properties designed for the NIAID research community: `funding, infectiousAgent, healthCondition`, and `species`. To improve consistency within categorical metadata properties, we used the same default ontologies for `infectiousAgent, healthCondition` and `species` across ComputationalTool and Dataset. A biomedically-relevant default ontology was inherited from Bioschemas for the featureList and applicationSubCategory properties, and biological ontologies were tied to the `measurementTechnique` property of the Dataset and ComptuationalTool schemas (*see Methods)*. Similar to our Dataset schema, the ComputationalTool schema is intentionally minimalistic; however, as it inherits features from the Bioschemas:ComputationalTool and Schema.org:SoftwareApplication classes, properties like `input` (defining the data inputs to the tool) could also be included, depending on the application.

**Table 2.** Comparison of three ComputationalTool schema specifications: Schema.org, Bioschemas, and the infectious disease-specific NIAID SysBio ComputationalTool. Properties required in the schema are denoted by (*). Note that Schema.org has additional properties not included in either Bioschemas or NIAID SysBio schemas which are omitted for simplicity.

| property | schema.org | Bioschemas | NIAID SysBio |
| --- | --- | --- | --- |
| applicationCategory | Text, URL | Text | Text |
| applicationSubCategory | Text, URL | Text, URL | Text, DefinedTerm |
| applicationSuite | Text | Text | NA |
| author | Organization, Person | Organization, Person | NA |
| citation | CreativeWork, Text | CreativeWork, Text | ScholarlyArticle |
| citation.name | Text | NA | Text |
| ↳ citation.url | URL | NA | Text* |
| codeRepository | URL | NA | URL |
| contributor | Organization, Person | Organization, Person | NA |
| creator | Organization, Person | Organization, Person | Organization, Person* |
| ↳ creator.name | Text | NA | Text* |
| dateModified | Date, DateTime | NA | Text |
| description | Text | Text* | Text* |
| discussionUrl | URL | URL | NA |
| downloadUrl | URL | URL* | NA |
| featureList | Text, URL | Text, URL | Text, DefinedTerm |
| funder | Organization, Person | Organization, Person | (see funding) |
| funding | Grant | NA | MonetaryGrant* |
| ↳ funding.description | NA | NA | Text |
| ↳ funding.funder | NA | NA | Organization* |
| ↳ funding.funder.name | NA | NA | Text* |
| ↳ funding.funder.parentOrganization | NA | NA | Text |
| ↳ funding.identifier | NA | NA | Text* |
| ↳ funding.url | NA | NA | Text |
| hasPart | CreativeWork | Dataset, URL | NA |
| healthCondition | MedicalCondition | NA | Text, DefinedTerm |
| identifier | PropertyValue, Text, URL | PropertyValue, Text, URL | Text* |
| infectiousAgent | Text | NA | Text, DefinedTerm |
| input | NA | FormalParameter | NA |
| isAccessibleForFree | Boolean | Boolean | NA |
| isBasedOn | CreativeWork, Product, URL | CreativeWork, Product, URL | NA |
| isPartOf | CreativeWork, URL | CreativeWork | NA |
| keywords | Text, URL | Text | NA |
| license | CreativeWork, URL | CreativeWork, URL | Text |
| name | Text | Text* | Text* |
| operatingSystem | Text | Text | NA |
| output | NA | FormalParameter | NA |
| programmingLanguage | ComputerLanguage, Text | NA | Text |
| provider | Organization, Person | Organization | NA |
| sameAs | URL | URL | NA |
| sdPublisher | Organization, Person | NA | Organization, Person |
| softwareAddOn | SoftwareApplication | SoftwareApplication | NA |
| softwareHelp | CreativeWork | CreativeWork | Creative Work |
| softwareVersion | Text | Text | Text |
| species | NA | NA | Text, DefinedTerm |
| thumbnailUrl | URL | URL | NA |
| url | URL | URL* | URL* |

### Encouraging metadata standardization, deposition, and registration through the Data Discovery Engine

After creating schemas to capture metadata for infectious disease datasets and computational tools, the next challenge is collecting metadata which conforms to these standards. One key barrier to using community standards like Schema.org for the deposition of data in a repository is identifying which of the 200+ available repositories provide metadata in the most discoverable format. To facilitate this process, we created a dataset registration flowchart (**Figure 2a**) based on our landscape analysis of biological repositories (**Supplemental Table 1**) that advises data providers how to register their dataset with minimal effort. If the dataset is stored in a Schema.org-compliant repository, the registered metadata can automatically be ingested via pre-existing markup crawlers like Google Dataset Search. Should the repository not be Schema.org-compliant, the data provider can register the deposited dataset by submitting the appropriate metadata via the NIAID SysBio Dataset Guide, as described below. To help users determine whether or not a data repository is Schema.org-compliant, we also built a web-based data portal/repository compatibility checker (**Figure 2b**). The data portal compatibility checker includes the more than 200 data repositories and portals commonly used by the biomedical research community we analyzed in our landscape analysis (**Supplemental Table 1**).

**Figure 2.**
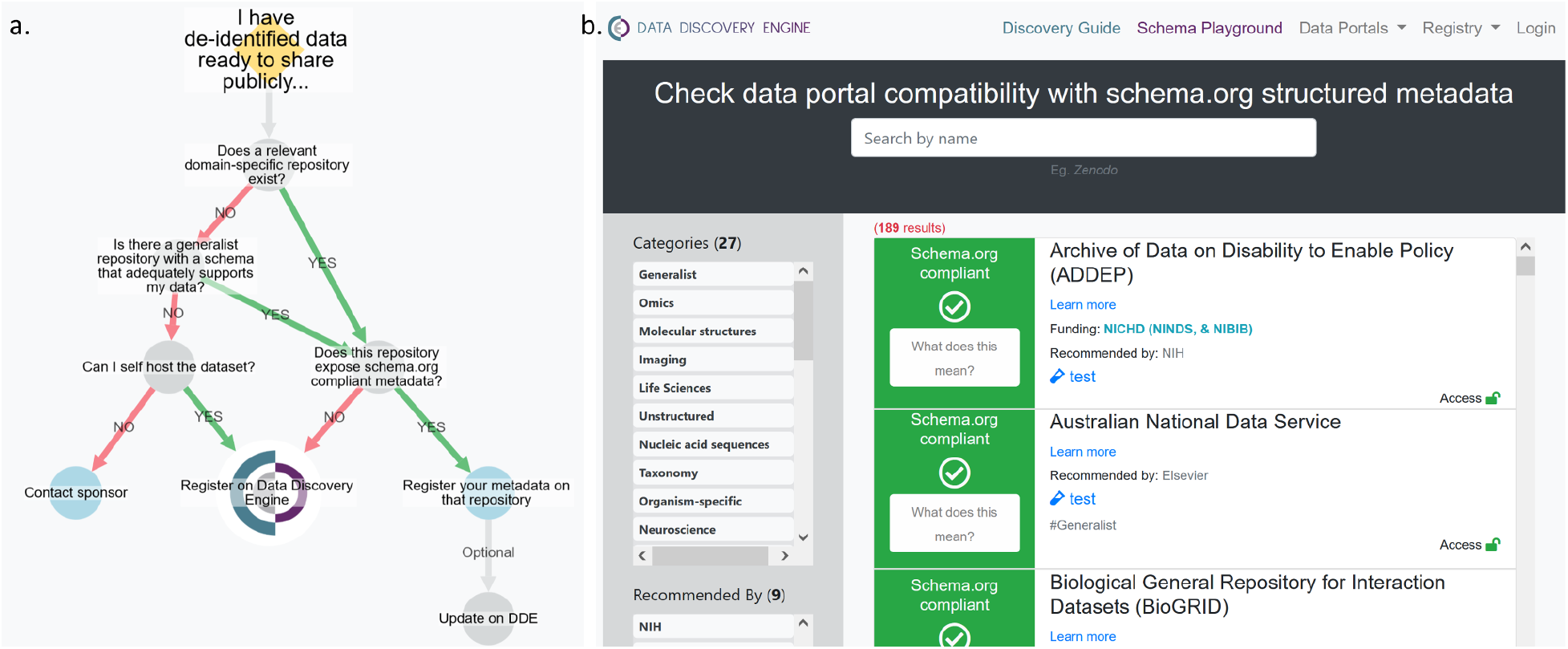
**a**, The DDE Dataset registration flowchart illustrates how to register a dataset with minimal effort. **b**, The portal compatibility checker tool helps identify Schema.org-compliant repositories. Available at https://discovery.biothings.io/compatibility.

For datasets deposited in repositories that are not Schema.org-compliant, or for data providers who want to add additional infectious disease-specific metadata, we leveraged the infrastructure of the Data Discovery Engine (DDE)^43^ to allow users to register their own metadata. The Discovery Guide creates easy-to-use forms which can be filled out manually or via imports from files stored on GitHub or on a local machine. These forms validate the input to ensure compliance with the schema and controlled vocabularies, improving the standardization and findability of the inputted metadata. The resulting pages^44–50^ for the registered datasets and tools contain embedded structured metadata, which automatically allow projects like Google Dataset Search to identify and index them.

The DDE also contains tools to improve ease of creation, use, and adoption of the schema. To enable others to reuse or extend our schemas^51^, we created and registered them using the DDE’s Schema Playground. Developing a schema is often a time-consuming process; the DDE contains tools which allow you to develop a schema in a Schema.org-compliant JSON for Linking Data (JSON-LD) graph format and share the resulting schema with others. Additionally, the Schema Playground includes tools allowing resource providers to easily extend and customize an existing schema based on a preexisting schema (*e.g*. Schema.org Dataset or NIAID SysBio ComputationalTool) to suit their needs. Our minimalistic NIAID SysBio Dataset schema is designed to be customized for particular use cases. For instance, in the Center for Viral Systems Biology Data Portal^52^, which focuses on collecting and publishing systems biology data from Lassa Fever, Ebola, and COVID-19 patients, we extended the NIAID SysBio Dataset schema to include additional properties to track within the consortium, including measurementCategory, publisher, and variableMeasured for specific enumerated values^53^. The tools within the DDE facilitate finding existing schemas and adapting them for new purposes, lowering the barrier to adoption and discouraging the creation of yet another bespoke schema which is incompatible with commonly used schemas.

### Applying the schemas to build a catalog of NIAID Systems Biology datasets & software

After creating a schema tailored to infectious and immunological data, we used the Data Discovery Engine infrastructure to register datasets created by the NIAID/DMID Systems Biology Consortium for Infectious Diseases (Systems Biology^54^). While the Systems Biology program has been prolific in generating publicly available datasets and open source computational tools, these research assets are challenging to find, as they are hosted in a combination of diverse data repositories, including specialist repositories, generalist, and custom data portals. As a result, accessing these datasets and computational tools becomes extraordinarily difficult. To address this limitation, the Systems Biology Consortium members across 15 different Centers individually registered 345 Datasets^55^ and 49 ComputationalTools^56^ in the DDE as of July 2022 (**Figure 3**). The datasets, encompassing host and pathogen relationships, come from over 18 diverse data repositories, including specialist repositories, generalist, and custom data portals. These datasets also span a range of experimental techniques, from omics (genomics, metagenomics, transcriptomics, lipidomics, proteomics) to immunological (flow cytometry, Systems Serology) to clinical. Importantly, we were cognizant of the potential burden on researchers to collect this information and sought to design an interface which minimized effort. All researchers we surveyed reported that the form-based submission made data registration easy and took on the order of a few minutes for each submission.

**Figure 3.**
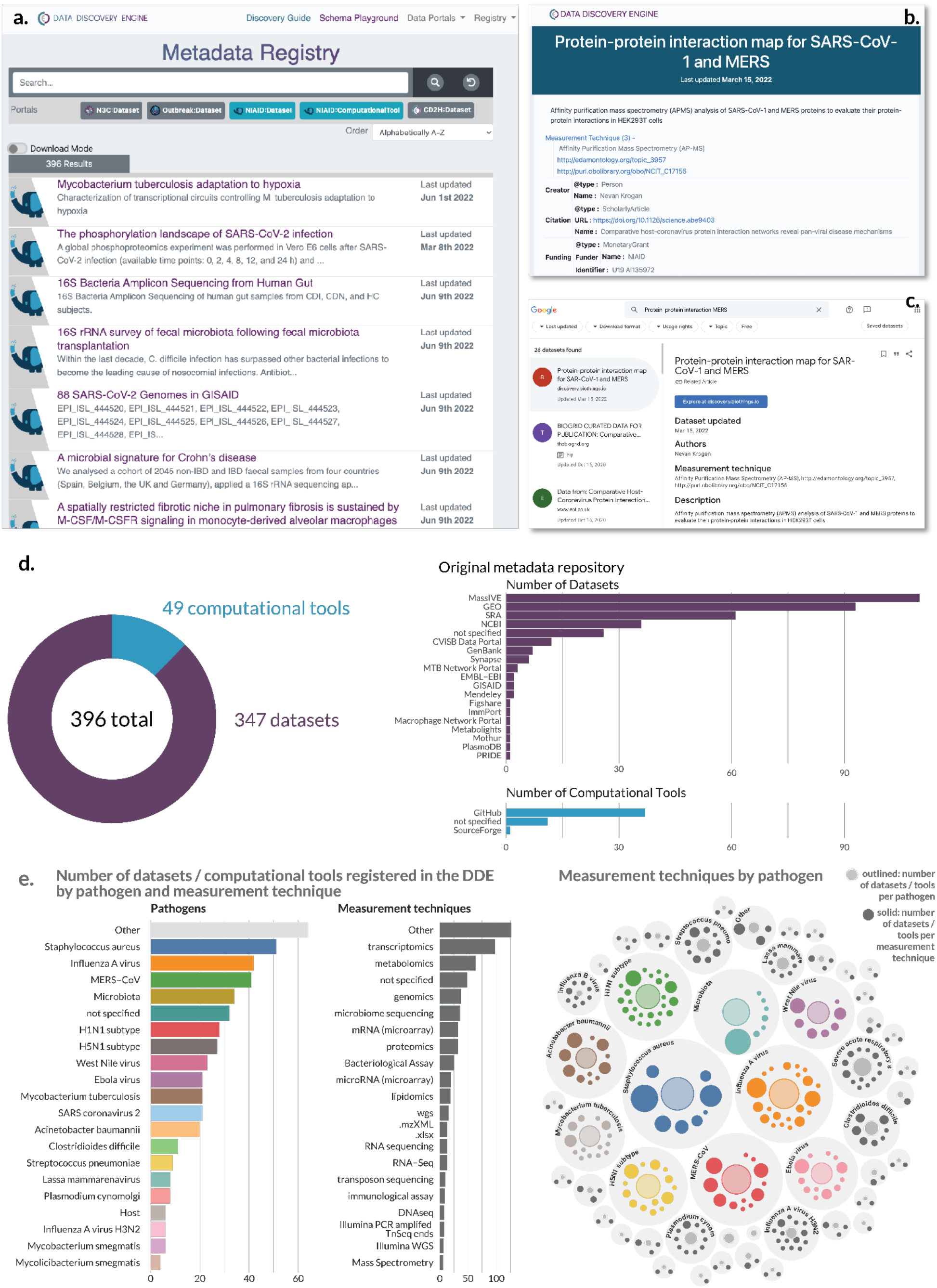
NIAID Systems Biology Consortium Dataset and ComputationalTool Catalog. The metadata is registered and available through the Data Discovery Engine (DDE). **a**, The DDE provides an interface to search for Datasets and ComputationalTools registered according to the NIAID SysBio schemas. **b**, Example metadata page^57^ for a Dataset registered on the DDE according to the NIAID SysBio schema. **c**, The same dataset in **Figure 3b** viewed in Google Dataset Search after its registration in the DDE. The standardized metadata is exposed as structured data markup, allowing web crawlers such as Google Dataset Search to discover them, increasing their findability. **c**, Summary statistics for the Datasets and ComputationalTools registered by the Systems Biology groups. **d**, Comparison of measurement techniques by pathogen in registered datasets.

Building a registry that allows the adoption of a common data schema to existing datasets provides a number of benefits to the data producer, data consumer, and research community at large. First, rapid data dissemination in this format increases the likelihood of secondary analyses on these datasets by others. Moreover, it has the potential to increase the number of citations to the dataset, benefiting both the user and the broader scientific community. For example, datasets registered on the DDE have their metadata exposed in a common format, making them findable via Google Dataset Search (**Figure 3c**) or other metadata harvesters. Second, data registry allows for the tracking of publicly available data directly by project sponsors, which would otherwise be difficult since data deposition often does not require the project funding as required metadata. Third, the metadata generated using this form has increased standardization along three axes: enforcing the collection of required information, enforcing consistent structure and format of fields, and ensuring consistency of terms via controlled vocabularies or ontologies for `measurementTechnique, species, infectiousAgent, and healthCondition`. For instance, the `healthCondition` field for COVID-19 has a number of synonyms or different spellings (*e.g*. COVID-19, Covid, Coronavirus disease 2019, nCoV-19); in the DDE, these are all standardized to a single value tied to an entry in the Mondo Disease Ontology^58,59^ (COVID-19, MONDO_0100096). This standardization should streamline data discovery and consequently promote dataset reuse. However, it is important to note that this effort represents a use case at this time, and the full impact of this standardized research catalog has yet to be realized.

### Use case: How a standardized registry benefits NIAID program sponsors

The registry generated by the Systems Biology Consortium (**Figure 3**) is currently being used to directly track and communicate the benefit of the research program by the NIAID sponsors. NIAID is committed to ensuring widespread access to all digital assets (publications, data, and analysis tools) that result from experimental approaches to exploit their full value and incentivize novel discoveries. While processes are established to track publications associated with specific grants and contracts (Research Performance Progress Reports, RPPR, which feeds into NIH RePORTER and MEDLINE / PubMed Central), datasets often require self-reporting from scientists to NIAID. In an ideal world, this information would be pulled automatically from repositories and/or PubMed, similar to publications; however, repositories do not always directly cite funding sources and/or critical metadata linking datasets to publications (and thus funding sources) are missing. Self-reporting limits the FAIRness of datasets by placing the reporting burden on the grantees, resulting in static snapshots of dataset availability which requires continuous updating. Additionally, in many instances, these reports are not visible to the scientific community, curtailing the ability of researchers to find and use these datasets in their research. As NIH moves towards its new data sharing policy in Fiscal Year 2023^7^, the Systems Biology Program, through its generation of large datasets and tools, serves as a model for data sharing best practices.

The NIAID SysBio Dataset schema enables landscape analysis of infectious disease and immunological datasets for both science administrators (*i.e.-* program officials) and the scientific community (*i.e.-* researchers). For a specific dataset, the DDE highlights the pathogen, associated experimental techniques, corresponding publications, repository location, and funding information such as grant and/or contract number (**Table 3**). The DDE allows program sponsors to evaluate datasets based on pathogen, linking pathogen-specific datasets, experimental techniques, and repositories. The DDE also connects experimental techniques (genomics, transcriptomics, proteomics, lipidomics, etc.) to specific pathogens enabling one to determine whether or not datatypes exist for particular pathogens (**Figure 3d**). Using the data collected through the registry, program sponsors can analyze the open data and open source tool contributions of research outputs from their program as a potential metric to measure program impact. It is important to note, however, that the scale of a dataset varies: a dataset can represent only a few samples/patients, or it could encompass tens of thousands of samples or more. As a result, the number of datasets or computational tools is merely the starting point to describe and evaluate the outputs of a research program. Taken together, the DDE harmonizes disparate datasets across repositories informing the community and program sponsors about infectious disease datasets.

**Table 3.** Overview of 345 datasets available in the DDE (as of 12 July 2022). Datasets are grouped by grant number and derived from multiple iterations of the Systems Biology Program for Infectious Diseases. * AI-11-038 (Contract^60^); ** AI-12-027 (U01/U19 grants^61^); *** AI-14-064 (U01 grants^62^); **** AI-16-080 (U19 grants^63^). Code to dynamically generate this table is available at https://github.com/Hughes-Lab/niaid-schema-publication/blob/main/figures-code/Table%203%20-%20Datasets%20by%20Grant.qmd.

| funding.identifier | author.name | infectiousAgent | MeasurementTechnique | Data Repositories |
| --- | --- | --- | --- | --- |
| HHSN272201200031C1* | MaPHIC | *Plasmodium cynomolgi* | Animal Data, Genomics, Immunological Assay, Lipidomics, Metabolomics, Proteomics | PlasmoDB, GEO, ImmPort, MassIVE, Metabolights, PRIDE |
| U01AI111598** | FluDyNeMo | Influenza A, Human, H1N1 | Immunological Assay, Microbiome Sequencing, Pathology, Transcriptomics | Synapse |
| U19AI106754** | FluOMICS | Influenza A, Avian (Including H5N1), Influenza A, Human, H1N1, Influenza A, Human, H3N2 | Genomics, Metabolomics, Proteomics, Software, Transcriptomics | GEO, MassIVE, Figshare, PI Website |
| U19AI106761** | Omics4TB | *Mycobacterium tuberculosis*, Other | Genomics, Software, Transcriptomics | MTB Network Portal, Macrophage |
|  |  |  |  | Network Portal,<br>GEO |
| U19AI106772** | OmicsLHV | Influenza A, Avian (Including H5N1), MERS-CoV, Ebola Virus, Influenza A, Human, H1N1, West Nile Virus, Other | Genomics, Lipidomics, Metabolomics, Proteomics, Transcriptomics | GEO, MassIVE |
| U01AI124255*** | Systems biology of *Clostridium difficile* infection | *Clostridium difficile*; Gram-Positive Bacteria; Microbiome/Microbiota | Software, Genomics, Transcriptomics, Metagenomics, Microbiome Sequencing | SRA |
| U01AI124275*** | Systems Biology of Microbiome-mediated Resilience to Antibiotic-resistant Pathogens | Microbiome/Microbiota | Metadata, Metagenomics, Microbiome Sequencing | Figshare, SRA |
| U01AI124290*** | Decoding Antibiotic-induced Susceptibility to *Clostridium difficile* Infection | *Clostridium difficile*; Microbiome/Microbiota | Genomics, Transcriptomics, Microbiome Sequencing | NCBI, SRA, Mothur |
| U01AI124302*** | Predicting the emergence of antibiotic resistance through multi-omics approaches and Immune System-surveillance | *Acinetobacter*, *Escherichia coli*, *Klebsiella*, *Pseudomonas aeruginosa*, *Salmonella*, *Staphylococcus aureus*, *Streptococcus pneumoniae* | Transposon Sequencing, Genomics, Transcriptomics, Software | SRA, NCBI |
| U01AI124316*** | Systems Biology Approach to Redefine Susceptibility Testing and Treatment of MDR Pathogens in the Context of Host Immunity | *Acinetobacter*, *Enterococcus*, *Escherichia coli*, Gram-Negative Bacteria, *Pseudomonas aeruginosa*, *Salmonella*, *Staphylococcus aureus* | *Acinetobacter*, Bacteriology Assay, Genomics, Metabolomics, Transcriptomics | MassIVE, SRA |
| U01AI124319*** | Systems Immunobiology of Antibiotic-Persistent MRSA Infection | *Staphylococcus aureus* | Reduced-Representation Bisulfite Sequencing | GEO |
| U19AI135972**** | FluOMICS: The Next Generation | Influenza A, Avian (Including H5N1); Influenza A, Human, H1N1; SARS-CoV-2 | Transcriptomics; Software | GEO, PI Website |
| U19AI135976**** | Omics4TB | *Mycobacterium tuberculosis*; *Mycobacterium*, Other | Genomics; Metabolomics; Transcriptomics; Software | SRA, GEO, NCBI |
| U19AI135990**** | HPMI | *Mycobacterium tuberculosis*; SARS-CoV-2 | Proteomics; Software; Crispr Ko Screens | Pride, NDEx, EMBD, Mendeley, Cytoscape |
| U19AI135995**** | CViSB | Ebola Virus; Lassa Fever Virus; SARS-CoV-2 | Genomics; Immunological Assay; Metadata; Virology Assay; Software | CViSB Website, Publication |
| U19AI135964**** | SCRIPT | *Acinetobacter*; Gram-Positive Bacteria; *Pseudomonas aeruginosa*; Microbiome/Microbiota; SARS-CoV-2 | Genomics; Metabolomics; Transcriptomics; Software; Metagenomics; Microbiome Sequencing | GenBank, Figshare, SRA, NCBI, GISAID, PI Website, GEO, Manuscript, Publication, Website |

### Use case: Adapting the NIAID SysBio Dataset and ComputationalTool schema to track COVID-19 research via the outbreak.info Research Library

One primary advantage of the minimalistic NIAID SysBio Dataset and ComputationalTool schemas is the ease with which they can be adapted and extended. By making the schemas available within the DDE, the time to develop a schema is lowered (reusability), the schemas are easier to combine with each other (interoperability), and the resulting metadata are easier to search or develop tools (findability). At the start of the COVID-19 pandemic, we noticed that an unprecedented number of research groups were sharing their research (publications, datasets, computational tools, and more) in near real-time, with thousands of items being shared every week^64^. While this shared knowledge has the potential to accelerate the pace of research, keeping track of the changing landscape is a challenge, with datasets, publications, and tools being published in dozens of different repositories, each with its own data schemas.

To address this challenge, we built the outbreak.info Research Library^64^, a standardized searchable platform of COVID-19 research. The Research Library harvests metadata from 16 sources every day and standardizes them to a schema^65^ based on the NIAID SysBio schemas, enabling querying across data platforms and across resource types. For instance, searching for “Omicron” allows researchers to simultaneously find publications, preprints, clinical trials, datasets, and websites related to the Variant of Concern without having to search the individual data sources. In this emerging health crisis, time was of the essence to develop a tool that could unify metadata from different sources. By using the NIAID SysBio schemas as our base, we accelerated the development of the system to enable rapid prototyping, testing, and deployment. Moreover, since the schemas share common parent classes, they naturally can be combined with data harvested according to the NIAID SysBio schemas, such as the Systems Biology Datasets and ComputationalTools. The flexibility of the system also allowed us to add additional properties customized to the goals of the Research Library. For instance, we added a property called `topicCategory`, based on machine learning on the metadata `name` and `description`, to group the datasets and publications into topics to support exploration of COVID-19-specific topics such as “Epidemiology” or “Public Health Interventions”. The outbreak.info Research Library has been accessed over 40,000 times in the past year from approximately 10,000 users in over 140 countries, demonstrating the community interest in searching standardized research metadata as well as the reach of projects which use or extend the NIAID SysBio schema.

Similarly, the NIAID SysBio Dataset and ComputationalTool schemas are also being used as the backbone in the development of the NIAID Data Ecosystem, a platform developed by NIAID to help immune-mediated and infectious disease researchers to discover and analyze data from diverse data sources. As more data projects adopt the core base of the NIAID SysBio and Schema.org schemas, their interoperability will increase and it will be easier to combine data from across sources.

## Discussion

The primary barriers to widely sharing and reusing data are a set of interrelated challenges (see companion Commentary on Addressing barriers in FAIR data practices for biomedical data ^66^), including: (***1***) lack of incentives for researchers to share information in a timely and transparent manner; (***2***) logistical challenges with collecting, sharing, and finding data; and (***3***) lack of documentation and interoperability conflicts which make it difficult to reuse data. As a result, our main goal in this work was to tackle the logistical barrier: collecting sufficient metadata that promotes easily discovering datasets, the first step in reusing public data.

Within the vast history of dataset sharing efforts, countless schemas have been developed and continue to be created. Previous efforts to improve the findability of disparate biomedical resources typically focused on developing exhaustive but often difficult-to-implement schemas.^67^ These approaches often aggregated a specific type of research outputs such as Datasets^68,69^ or Experiments^70^, or applying Natural Language Processing (NLP) algorithms to improve the searchability of insufficiently described research outputs^71–74^. For example, DataMed ^68,69^, a biomedical research dataset index funded through the Big Data to Knowledge initiative, indexed and mapped metadata to core Data Tag Suite (DATS) elements, conducted user experience studies^75^, and applied NLP algorithms to improve the searchability of the indexed Datasets^69^. Despite the solidity of this approach, however, there is limited evidence of uptake of their schema, possibly due to the barriers in sharing data and reusing metadata schemas. As a result, the DataMed site is still a beta version and the metadata has not been updated since 2017.

Our goal in creating the NIAID SysBio Dataset and ComputationalTool schemas was not to create yet *another* schema (which may or may not be widely adopted). Instead, we sought to develop a process, reusable across diverse areas of biomedical investigation, to create a schema which is interoperable with community projects but tailored for a specific purpose. Given the diversity of data that exists, it is unlikely that the community will ever settle on a single schema; as a result, our approach focuses on highlighting the core properties shared between schemas and developing crosswalks to translate between schemas. In this respect, we are following a model similar to the infrastructure of PubMed, where each journal can develop its own data model, but must contain a common set of properties to be findable within PubMed.

An overarching challenge to meaningful data dissemination is the lack of a core set of properties for describing data, contributing to the fragmented data landscape and making data hard to find. In our survey of over 200 generalist and biological repositories, only 18% of biological repositories used a common standard to describe their metadata, the widely adopted Schema.org Dataset standard (**Figure 1**). As a result, data consumers are forced to search for different information in each data source, making datasets more challenging to find. Furthermore, cross-repository searching projects like Google Dataset Search or the NIAID Data Ecosystem are often forced to either undergo time-consuming metadata standardization projects or ignore potentially valuable metadata that cannot easily be standardized.

Based on this survey of existing dataset metadata schema usage, we developed a minimalistic, extendable schema tailored to infectious and immune-mediated disease datasets/software (**Tables 1 and 2**). To promote interoperability, we chose to base our schema on Schema.org to ensure compatibility with data cataloging projects like Google Dataset Search and BioSchemas. As our goal was not to create a new standard, but to generate links between standards, our approach was to extend existing classes with new properties to promote findability of infectious and immune-mediated disease assets. Essential to promote translation between standards, we developed a crosswalk to other common schemas, based on the work of the Research Metadata Schemas Working Group of the Research Data Alliance^42^ (see **Supplemental Table 2**).

A core driver of our approach for developing a standard was to focus on practicality to decrease the barrier for research groups to collect metadata. While exhaustive schemas provide a full set of information about a dataset or computational tool, in practice, they can be inefficient to use. Indeed, if a Schema.org Dataset used every available property, a research group would need to capture over 124 pieces of information – a time consuming task, particularly when many of the properties are irrelevant for biological datasets. By collecting a small, core set of metadata properties, we focused data collection efforts – which can be quite time consuming – on essential fields to capture the gestalt of the dataset. We identified the minimum amount of information needed to understand what a dataset contains based first on understanding what Dataset properties are commonly used across dataset providers (**Figure 1**). Such properties were then supplemented with a relatively small number of infectious and immune-mediated disease-specific fields based on discussions within the Systems Biology Data Dissemination working group, which contains experimentalists, computational biologists, clinicians, and NIAID program staff. Additionally, to decrease effort in post-processing of metadata, such as the efforts to use Natural Language Processing to improve metadata FAIRness by categorizing and classifying unstructured text^69,72^, we leverage existing key ontologies for properties to minimize variability in categorical metadata values.

To test how this schema could be deployed in a real-world situation, we leveraged the 15 Centers within the NIAID Systems Biology Consortium for Infectious Diseases, representing research groups from over 30 different institutions, to register Datasets and ComputationalTools created by their center. Using the Data Discovery Engine (DDE) registry guide, the centers registered nearly 400 datasets and tools, which are physically stored in over 18 different sources (**Figure 3**). The resulting registry^55,56^, therefore, provides a single searchable interface of the research output of the Systems Biology program and exposes its structured metadata for data indexing projects such as Google Dataset Search to find. As the DDE registration enforces type, metadata structure, and links to ontologies or controlled vocabularies, the metadata is more easily searched. For example, the standardized, centralized metadata enables funding agencies to track research outputs and monitor sharing compliance. Such transparency will be essential as the NIH launches its new Data Sharing Policy^7^. A centralized metadata registry also decreases the burden of manual data reporting to funding agencies and enables funding agencies to rapidly respond to congressional requests.

Our approach of creating a common schema with core interoperable elements that can be extended to particular use cases has also been replicated in other projects, increasing the opportunities to promote cross-resource searching. Already, the NIAID SysBio Dataset and ComputationalTool schemas have been used in a variety of other infectious disease data sharing projects, including a consortium-specific Data Portal (Center for Viral Systems Biology Data Portal^52,53^), a searchable COVID-19 research platform (outbreak.info Research Library^64^), and an immune-mediated and infectious disease Discovery Portal (NIAID Data Ecosystem).

Standardizing metadata properties is an important first step for improving data FAIRness, but does not solve all problems associated with making data findable, accessible, interoperable, and reusable. In particular, organizing, managing, combining, and interpreting large datasets – especially across different research areas and groups – remain core challenges. While artificial intelligence and machine learning may provide avenues to address some of these issues^73^, those methods rely on having sufficient and unbiased data to supply as inputs, necessitating researchers to invest in collecting sufficient metadata and data. Our work here aims to lower the barriers to register datasets with a core set of metadata to address challenges in findability, which will then stimulate iterative improvements on the downstream aspects of data sharing, including data interoperability.

We hypothesize that these two factors – a user friendly minimalistic dataset registration and the enforced standardization of the metadata using a schema containing a common core of elements – will promote data dissemination, improved findability of datasets (eventually leading to increased dataset citations), and acceleration of research. We further hypothesize that this process is self-reinforcing: as data sharing and reuse becomes easier, the incentives to provide better quality metadata and data will increase, driven by use cases and reuse of data (quantified through citations). In this manner, metadata registration of datasets scattered across repositories is analogous to Open Access research articles, which have been shown to be more immediately recognized and cited by peers than non-Open Access articles, even if that journal is widely available^76^. As such, the registry represents a step towards promoting a cultural shift to rapid data dissemination and findability. By lowering the barrier to finding data, the metadata registry has the potential to motivate scientists to share data in repositories as data digital object identifiers (DOI) can be cited during secondary data analysis. This strategy also enables funding agencies to identify (and potentially reward) grants and grantees that represent exceptional data sharing best practices. Ensuring that datasets are disseminated following the FAIR Data Principles represents a challenge beyond data collection and analysis, and is essential to ensure that publicly available data is reproducible and interpretable for secondary analyses by investigators who did not generate the data. As has been abundantly evidenced by the necessity to readily share and analyze large datasets arising during the COVID-19 pandemic, integrative sharing of complex and diverse datasets is imperative to advance research focusing on human health. The effort from this Working Group has started to address these challenges and demonstrates “big data” can be incorporated across research programs, will generate meaningful results, and will continue to be an integral component of research activities across infectious diseases. Taken together, FAIR data, analytical software, computational models, protocols, and reagents represent invaluable resources for the infectious disease community as we continue to understand and respond to the impacts of existing and emerging pathogens on human disease.

## Conclusions

We have developed a reusable process for improving the FAIRness of scientific research outputs which leverages the flexibility of Schema.org while being adaptive to the needs of specific research communities. Our goal was not to create yet another bespoke schema which may or may not be adopted, but to focus on developing a process to create a schema which can be easily integrated with metadata based on other common schemas. To achieve this goal, we evaluated Dataset repositories for Schema.org compliance, created interoperable Dataset and ComputationalTool schemas that considers the needs of the Systems Biology research consortia and its stakeholders, and identified additional challenges to FAIRness that have yet to be addressed. We leveraged existing open source infrastructure and utilized our process to create a data portal which improves FAIRness of Allergy and Infectious Disease datasets and tools. In doing so, we lower the barrier to reuse of datasets and tools in this research space, enable meta-analyses in this area, and provide a reusable process and working example for improving data FAIRness to other research communities.

## Methods

### Survey of Schema.org adoption across repositories

We undertook a survey of biomedical repositories to try to understand current usage of the Schema.org Dataset schema across generalist and biologically-focused sources. To begin, we identified over two hundred repositories recommended by NIH^22^, other donor agencies^9,10,77^, journals^3–6,78–80^, and on FAIRSharing.org^81^. We then analyzed whether a randomly selected dataset from each repository contained Dataset metadata markup that was Schema.org-compliant using Google’s Rich Results Test Tool (https://search.google.com/test/rich-results). For a subset of these repositories, we harvested all Dataset metadata available from 7 sources using custom metadata harvesting infrastructure. This custom metadata harvesting infrastructure consists of an extendable metadata crawler that could be used seamlessly within the BioThings^82^ and Data Discovery Engine^43^ frameworks. The crawler consists of three primary components which include source-specific spiders, analytical scripts, and uploaders that help load the resulting crawled metadata into a BioThings-based API^82^. The source-specific scrapers act like BioThings plugins for a particular source or repository of interest. If a sitemap is already available, the sitemap scraper ingests the sitemap URL, the sitemap rules, and any recommended user agents to systematically crawl through and apply the content scraper to parse out Schema.org-compliant metadata. If a sitemap is unavailable, the content scraper needs to include helper functions for calling the API (if the resource provides its metadata via API), paginating through a source to crawl and parse each page, or render and parse dynamically generated pages. Once a spider has been generated for each source, a corresponding uploader will be needed to map the results from the spider to the Elasticsearch index to enable querying of the metadata from a BioThings-based API. The metadata crawler is built in Python using the Scrapy library and can be found at https://github.com/biothings/biothings.crawler.

### Converting survey results to overall schema design decisions

As part of the NIAID Systems Biology Data Dissemination Working group, we identified the minimal metadata fields needed to describe a resource within a dataset or software class based on our survey of utilization of Schema.org Dataset properties (**Figure 1**), comparison to other commonly used community standards like Bioschemas (**Table 1 and Extended Table 2**), and our own data collection and reuse experience. The NIAID/DMID Systems Biology Consortium for Infectious Diseases is a group of interdisciplinary scientists that bridge disparate scientific disciplines including microbiology, immunology, infectious diseases, microbiome, mathematics, physics, bioinformatics, computational biology, machine learning, statistical methods, and mathematical modeling. These teams work together to integrate large-scale experimental biological and clinical data across temporal and spatial scales through computational and modeling tools to better understand infectious diseases. The research findings drive innovation and discovery, with the goal of developing novel therapeutic and diagnostic strategies, and predictive signatures of disease to alleviate infectious disease burden and provide solutions to complex public health challenges and disease outbreaks. Within this Consortium, the Data Dissemination Working Group involves representatives from each of the Systems Biology Centers. The members of this group work together to establish best practices for disseminating systems biology data in a way that is consistent with the NIAID Data Management and Sharing Guidelines^83^ and with the FAIR principles^15^.

Based on our survey of utilization of Schema.org Dataset properties, we identified the minimal metadata fields needed to describe a resource within a dataset or software class. Using our survey, we calculated the prevalence of properties within a source and then obtained the average prevalence for each property across the seven sources surveyed. Properties with an average prevalence of greater than 70 percent were assigned as minimal. We then supplemented these core properties with additional properties prioritized for infectious disease discoverability based on our experience finding and reusing datasets and computational tools. Given the importance of funders as key stakeholders in data access, we included funding as a minimal property as well. Although `measurementTechnique` did not make the prevalence cutoff, it was determined to be invaluable towards future aspirations for linking datasets and computational tools, and developing integrated analytical pipelines. Given the importance of spatial/temporal data in the infectious disease research space, we included properties to structure and store this information, even if its inclusion in the schema is currently mostly aspirational. The properties included in our schemas are not intended to be a comprehensive set of properties that any biomedical dataset could have, but rather a prioritized set of fields to promote reuse in biomedical research. Additionally, since our schemas are derived from Schema.org and Bioschemas properties, additional properties from those schemas can be inherited in derived schemas.

Going beyond the Schema.org standard, we reviewed whether each property should be required, recommended, or optional (marginality), and whether the value of a property should be single or multiple (cardinality) to ensure consistency in the application of our schema. Fields are only required if absolutely necessary to minimize metadata collection time, and we were intentionally parsimonious about what we selected. To further improve consistency, interoperability, and linkage between the classes, we added validations for controlled vocabularies based on standard ontologies, including NCBI Taxonomy^84^, NCI Thesaurus^85^, EDAM - Bioscientific data analysis ontology^86^, and the Mondo Disease Ontology^58^.

### Cataloguing NIAID SysBio Datasets and ComputationalTools within the Data Discovery Engine

To register Dataset and ComputationalTool metadata, we used the Data Discovery Engine’s^43^ Metadata Registry Guide. The Guide (Datasets: https://discovery.biothings.io/guide/niaid, ComputationalTools: https://discovery.biothings.io/guide/niaid/ComputationalTool) enables users to submit metadata according to the NIAID SysBio schema and validate their inputs. Representatives from across the NIAID SysBio consortium collected metadata for datasets and computational tools they had generated and registered them on the DDE using a combination of form-based entry and .csv-based bulk uploads. The registered metadata can be viewed at https://discovery.biothings.io/dataset?guide=/guide/niaid,/guide/niaid/ComputationalTool.

## Data Availability Statement

The NIAID SysBio Dataset and ComputationalTool schemas are available in the Data Discovery Engine (https://discovery.biothings.io/view/niaid), and crosswalks between these schemas and other commonly used schemas are available in Supplemental Table 2. Schema.org-compliant repositories are available in Supplemental Table 1 and at https://docs.google.com/spreadsheets/d/1K1Jqy3CeCtRYtgC3RiC1FICqQFA8sCy2h5UafiAFk-U (Figure 1a). The harvested metadata from 6 selected repositories (Figure 1b) is available through the Metadata Crawler API (https://crawler.biothings.io, https://crawler.biothings.io/api/query https://crawler.biothings.io, https://crawler.biothings.io/api/query). NIAID Systems Biology Consortium Datasets and ComputationalTools registered on the Data Discovery Engine using the NIAID SysBio schemas are available on the DDE at https://discovery.biothings.io/dataset?guide=/guide/niaid,/guide/niaid/ComputationalTool and within the underlying API, https://discovery.biothings.io/api/dataset/query?q=_meta.guide:(“/guide/niaid”OR“/guide/niaid/ComputationalTool”)). The surveys have been archived as a Zenodo dataset for persistence at https://zenodo.org/record/7032896 (DOI 10.5281/zenodo.7032896).

## Code Availability Statement

Code used to analyze the prevalence of Schema.org properties and create figures is available at GitHub (https://github.com/Hughes-Lab/niaid-schema-publication and archived on Zenodo (DOI 10.5281/zenodo.6816052): https://zenodo.org/record/6816052. Code used to harvest metadata via Metadata Crawler (https://crawler.biothings.io/, https://github.com/biothings/biothings.crawler) and Metadata Plus (https://metadataplus.biothings.io/, https://github.com/biothings/metadataplus) are available on GitHub, and the code to create the Data Discovery Engine, including the NIAID SysBio Dataset and ComputationalTool registration guides, is available on GitHub (https://github.com/biothings/discovery-app). Code to dynamically generate Table 3 is available at https://github.com/Hughes-Lab/niaid-schema-publication/blob/main/figures-code/Table%203%20-%20Datasets%20by%20Grant.qmd.

## Acknowledgements

This work was supported in part by the National Institute of Allergy and Infectious Diseases (NIAID), National Institutes of Health (NIH) grants **U01 AI124290** (Baylor: TS, QW), **U01 AI124302** (Boston College: JB), **U19 AI135995** (Scripps Research: MAAC, LDH, GT, AIS, CW, JX, XZ), **U19 AI135964** (Northwestern: MK, LVR, JS), **U19 AI135972** (Sanford Burnham Prebys: LP); **U01 AI124319** (UCLA: MY), **75N91019D00024** (Scripps Research: CZ, GT, LDH, AIS, CW); National Center for Advancing Translational Sciences NIH grant **U24 TR002306** (Scripps Research: MAAC, LDH, GT, AIS, CW, JX, XZ); and National Institute of General Medical Sciences grant **R01 GM083924** (Scripps Research: MAAC, GT, AIS, CW, JX, XZ). We acknowledge the NIAID/DMID Systems Biology Consortium for Infectious Diseases Data Dissemination Working Group for developing the NIAID SysBio schemas, registering center-created datasets and computational tools, and providing critical feedback on the manuscript. We thank Reed Shabman for his leadership within the Data Dissemination Working Group, coordinating with centers to register datasets and tools, and helpful comments and careful revisions of the paper. We additionally thank Liliana Brown for the support of the Program this paper originated from and Serdar Turkarslan, Ishwar Chandramouliswaran, Wilbert van Panhuis, and Jack DiGiovanna for helpful discussions in preparing this manuscript.

## Author Contributions

**LDH, LP, RSS, AIS**, and **GT** created and/or provided feedback on the NIAID SysBio schema. **MAAC, CC, LDH, RSS, AIS, GT, CW, JX**, and **XZ** designed, developed, and provided feedback on the Data Discovery Engine metadata registration tool. **LDH, MK, LP, LVR, RSS, GT**, and **QW** compiled and/or registered metadata on the Data Discovery Engine guide. **JB, LB, LDH, MK, LP, TCS, LVR, JS, RSS, AIS, GT, CW, QW**, and **MRY** contributed to writing and editing the manuscript.

## NIAID Systems Biology Data Dissemination Working Group Members

Ginger Tsueng, José Bento, Lars Pache, Tor C. Savidge, Justin Starren, Luke V. Rasmussen, Mengjia (Marjorie) Kang, Qinglong Wu, Serdar Turkarslan, Michael R. Yeaman, Andrew I. Su, Chunlei Wu, Reed S. Shabman, Laura D. Hughes

## Bibliography

1. Siebert, M. et al. Data-sharing recommendations in biomedical journals and randomised controlled trials: an audit of journals following the ICMJE recommendations. BMJ Open 10, e038887 (2020).

2. Springer Nature Data Availability Statements. Springer Nature https://www.springernature.com/gp/authors/research-data-policy/data-availability-statements/12330880.

3. Science Data and Code Deposition Policy. Science Journals: editorial policies https://www.science.org/content/page/science-journals-editorial-policies.

4. The EMBO Journal: Author Guidelines. https://www.embopress.org/page/journal/14602075/authorguide doi:10.1002/(ISSN)1460-2075.

5. Information for Authors: Cell. https://www.cell.com/cell/authors.

6. PLOS ONE: Recommended Repositories. https://journals.plos.org/plosone/s/recommended-repositories.

7. NOT-OD-21-013: Final NIH Policy for Data Management and Sharing. https://grants.nih.gov/grants/guide/notice-files/NOT-OD-21-013.html.

8. Kozlov, M. NIH issues a seismic mandate: share data publicly. Nature Publishing Group UK http://dx.doi.org/10.1038/d41586-022-00402-1 (2022) doi:10.1038/d41586-022-00402-1.

9. Open Data at NSF. https://www.nsf.gov/data/.

10. Gates Open Research Data Guidelines. The Gates Forundation https://gatesopenresearch.org/for-authors/data-guidelines.

11. Wellcome Data, software and materials management and sharing policy. Wellcome Trust https://wellcome.org/grant-funding/guidance/data-software-materials-management-and-sharing-policy.

12. Errington, T. M., Denis, A., Perfito, N., Iorns, E. & Nosek, B. A. Challenges for assessing replicability in preclinical cancer biology. Elife 10, (2021).

13. Tedersoo, L. et al. Data sharing practices and data availability upon request differ across scientific disciplines. Sci Data 8, 192 (2021).

14. Gabelica, M., Bojčić, R. & Puljak, L. Many researchers were not compliant with their published data sharing statement: mixed-methods study. J. Clin. Epidemiol. (2022) doi:10.1016/j.jclinepi.2022.05.019.

15. Wilkinson, M. D. et al. The FAIR Guiding Principles for scientific data management and stewardship. Sci Data 3, 160018 (2016).

16. Musen, M. A. Without appropriate metadata, data-sharing mandates are pointless. Nature Publishing Group UK http://dx.doi.org/10.1038/d41586-022-02820-7 (2022) doi:10.1038/d41586-022-02820-7.

17. Howcroft, G. A Beginner’s Guide to Metadata and Keywords. Editors’ Bulletin 3, 75–77 (2007).

18. Ulrich, H. et al. Understanding the Nature of Metadata: Systematic Review. J. Med. Internet Res. 24, e25440 (2022).

19. Leipzig, J., Nüst, D., Hoyt, C. T., Ram, K. & Greenberg, J. The role of metadata in reproducible computational research. Patterns (N Y) 2, 100322 (2021).

20. Wilson, S. L. et al. Sharing biological data: why, when, and how. FEBS Lett. 595, 847–863 (2021).

21. NIH Scientific Data Sharing. https://sharing.nih.gov/.

22. NIH Data Sharing Resources. https://www.nlm.nih.gov/NIHbmic/nih_data_sharing_repositories.html (2013).

23. RDA COVID-19 Working Group. RDA COVID-19 Recommendations and Guidelines on Data Sharing. (2020).

24. Dugan, V. G. et al. Standardized metadata for human pathogen/vector genomic sequences. PLoS One 9, e99979 (2014).

25. Wei, W. et al. Finding relevant biomedical datasets: the UC San Diego solution for the bioCADDIE Retrieval Challenge. Database 2018, (2018).

26. Callaghan, S. Data Sharing in a Time of Pandemic. Patterns (N Y) 1, 100086 (2020).

27. Snijder, B., Kandasamy, R. K. & Superti-Furga, G. Toward effective sharing of high-dimensional immunology data. Nat. Biotechnol. 32, 755–759 (2014).

28. Foraker, R. E. et al. Transmission dynamics: Data sharing in the COVID-19 era. Learn Health Syst e10235 (2020) doi:10.1002/lrh2.10235.

29. Sansone, S.-A. et al. DATS, the data tag suite to enable discoverability of datasets. Sci Data 4, 170059 (2017).

30. Fenner, M. et al. A data citation roadmap for scholarly data repositories. Sci Data 6, 28 (2019).

31. Shepherd, A. et al. Science-on-Schema.org v1.3.0. (2022). doi:10.5281/zenodo.6502539.

32. Che, H. & Duan, Y. On the Logical Design of a Prototypical Data Lake System for Biological Resources. Front Bioeng Biotechnol 8, 553904 (2020).

33. Noy, N. Discovering millions of datasets on the web. Google: The Keyword (2020).

34. Facilitating the discovery of public datasets. Google AI Blog https://ai.googleblog.com/2017/01/facilitating-discovery-of-public.html (2017).

35. Benjelloun, O., Chen, S. & Noy, N. Google Dataset Search by the Numbers. arXiv[cs.IR] (2020).

36. Profiti, G. et al. Using community events to increase quality and adoption of standards: the case of Bioschemas. F1000Res. 7, (2018).

37. Michel, F. & The Bioschemas Community. Bioschemas & Schema.org: a Lightweight Semantic Layer for Life Sciences Websites. BISS 2, e25836 (2018).

38. Dataset Documentation for Google Dataset Search. Google Developers https://developers.google.com/search/docs/advanced/structured-data/dataset.

39. Bioschemas Dataset - 0.3 Release 2019_06_14. https://bioschemas.org/profiles/Dataset/0.3-RELEASE-2019_06_14.

40. King, G. An Introduction to the Dataverse Network as an Infrastructure for Data Sharing. Sociol. Methods Res. 36, 173–199 (2007).

41. International Food Policy Research Institute (IFPRI). COVID-19 Impact on Rural Men and Women in Ghana, Round 6. (2022) doi:10.7910/DVN/ZKGPQO.

42. Wu, M. et al. A Collection of Crosswalks from Fifteen Research Data Schemas to Schema.org. RDA https://www.rd-alliance.org/group/research-metadata-schemas-wg/outcomes/collection-crosswalks-fifteen-research-data-schemas (2021).

43. Cano, M. et al. Schema Playground: A tool for authoring, extending, and using metadata schemas to improve FAIRness of biomedical data. bioRxiv 2021.09.02.458726 (2022) doi:10.1101/2021.09.02.458726.

44. Viral Hemorrhagic Fever Consortium / Kenema Government Hospital. Blood Cell Counts of Ebola/Lassa Patients. Data Discovery Engine https://discovery.biothings.io/dataset/9f2318febbbfa710.

45. HPMI: Host Pathogen Mapping Initiative. Functional genomic screens to identify host factors for SARS-COV-2, OC43, and 229E. Data Discovery Engine https://discovery.biothings.io/dataset/60c702f2b5a0049d (2022).

46. University of Pittsburgh. Predicting the emergence of antibiotic resistance through multiomics approaches and Immune System-surveillance. Data Discovery Engine https://discovery.biothings.io/dataset/8a035090d274bf48.

47. Spinler, J., Savidge, T. & Baylor College of Medicine. C. difficile isolates from asymptomatic carriers and CDI patients. Data Discovery Engine https://discovery.biothings.io/dataset/758b3e902b1547e1.

48. Chang, Y.-L. & Lundquist Institute for Biomedical Innovation at Harbor-UCLA Medical Center. DNA methylation data from human patients infected with MRSA. Data Discovery Engine https://discovery.biothings.io/dataset/ea7518f519acc4b9.

49. Fluomics: The Next Generation. Role of diverse NS1 influenza segments in the infection of human bronchial epithelial cells. Data Discovery Engine https://discovery.biothings.io/dataset/bd813a34e9c9140d.

50. Successful Clinical Response In Pneumonia Therapy (SCRIPT) Systems Biology Center. Circuits between infected macrophages and T cells in SARS-CoV-2 pneumonia. Data Discovery Engine https://discovery.biothings.io/dataset/dc386eb3a37ba7a2.

51. Tsueng, G. et al. NIAID schemas. Data Discovery Engine https://discovery.biothings.io/portal/niaid.

52. CViSB Data Portal. Center for Viral Systems Biology https://cvisb.org/data/.

53. CViSB Schemas. Center for Viral Systems Biology Data Portal https://data.cvisb.org/schema.

54. Systems Biology Consortium for Infectious Diseases. https://www.niaid.nih.gov/research/systems-biology-consortium.

55. Systems Biology Datasets registered on the DDE. Data Discovery Engine https://discovery.biothings.io/dataset?guide=/guide/niaid.

56. Systems Biology ComputationalTools registered on the DDE. http://discovery.biothings.io/ https://discovery.biothings.io/dataset?guide=/guide/niaid/ComputationalTool.

57. Krogan, N. Protein-protein interaction map for SARS-CoV-1 and MERS. Data Discovery Engine https://discovery.biothings.io/dataset/e74bdfeef8542189.

58. Shefchek, K. A. et al. The Monarch Initiative in 2019: an integrative data and analytic platform connecting phenotypes to genotypes across species. Nucleic Acids Res. 48, D704–D715 (2020).

59. Mungall, C. J., Koehler, S., Robinson, P., Holmes, I. & Haendel, M. k-BOOM: A Bayesian approach to ontology structure inference, with applications in disease ontology construction. bioRxiv 048843 (2019) doi:10.1101/048843.

60. NOT-AI-11-038: RFP Announcement: An Integrated Approach to Understanding Host-Pathogens Interactions - RFP NIAID-DMID-NIHAI2010100. https://grants.nih.gov/grants/guide/notice-files/not-ai-11-038.html.

61. RFA-AI-12-027: OMICS Technologies For Predictive Modeling of Infectious Diseases (U19). https://grants.nih.gov/grants/guide/rfa-files/RFA-AI-12-027.html.

62. RFA-AI-14-064: Systems Biology and Antibacterial Resistance (U01). https://grants.nih.gov/grants/guide/rfa-files/rfa-ai-14-064.html.

63. RFA-AI-16-080: Systems Biology: The Next Generation for Infectious Diseases (U19). https://grants.nih.gov/grants/guide/rfa-files/rfa-ai-16-080.html.

64. Tsueng, G. et al. Outbreak.info: A standardized, searchable platform to discover and explore COVID-19 resources and data. bioRxiv 2022.01.20.477133 (2022) doi:10.1101/2022.01.20.477133.

65. Tsueng, G. et al. outbreak.info schemas. Data Discovery Engine https://discovery.biothings.io/view/outbreak.

66. Hughes, L. D. et al. Addressing barriers in FAIR data practices for biomedical data.https://docs.google.com/document/d/1w7zBq772fb5DUfbrdnTgZzcI1gVg3xFhlPk5EV54TPo/edit?usp=sharing.

67. Marc, D. T., Beattie, J., Herasevich, V., Gatewood, L. & Zhang, R. Assessing Metadata Quality of a Federally Sponsored Health Data Repository. AMIA Annu. Symp. Proc. 2016, 864–873 (2016).

68. Ohno-Machado, L. et al. Finding useful data across multiple biomedical data repositories using DataMed. Nat. Genet. 49, 816–819 (2017).

69. Chen, X. et al. DataMed - an open source discovery index for finding biomedical datasets. J.Am. Med. Inform. Assoc. 25, 300–308 (2018).

70. LINCS Phase II Extended Metadata Standards. NIH LINCS Program https://lincsproject.org/LINCS/data/standards.

71. Löbe, M., Stäubert, S., Goldberg, C., Haffner, I. & Winter, A. Towards Phenotyping of Clinical Trial Eligibility Criteria. Stud. Health Technol. Inform. 248, 293–299 (2018).

72. Wang, Y., Rastegar-Mojarad, M., Komandur-Elayavilli, R. & Liu, H. Leveraging word embeddings and medical entity extraction for biomedical dataset retrieval using unstructured texts. Database 2017, (2017).

73. Burgdorf, A., Pomp, A. & Meisen, T. Towards NLP-supported Semantic Data Management. arXiv [cs.IR] (2020).

74. Chung, G. Y.-C. Towards identifying intervention arms in randomized controlled trials:extracting coordinating constructions. J. Biomed. Inform. 42, 790–800 (2009).

75. Dixit, R. et al. User needs analysis and usability assessment of DataMed - a biomedical data discovery index. J. Am. Med. Inform. Assoc. 25, 337–344 (2018).

76. Eysenbach, G. Citation advantage of open access articles. PLoS Biol. 4, e157 (2006).

77. Wellcome Trust Data Guidelines. https://wellcomeopenresearch.org/for-authors/data-guidelines.

78. Nature Recommended Data Repositories. https://www.nature.com/sdata/policies/repositories.

79. Elsevier. Elsevier Database Linking. https://www.elsevier.com/authors/tools-and-resources/research-data/data-base-linking.

80. eLife Journal Policies. https://submit.elifesciences.org/html/elife_author_instructions.html#policies.

81. FAIRsharing Databases. https://fairsharing.org/databases/.

82. Lelong, S. et al. BioThings SDK: a toolkit for building high-performance data APIs in biomedical research. Bioinformatics (2022) doi:10.1093/bioinformatics/btac017.

83. Data Management and Sharing Guidelines. https://www.niaid.nih.gov/research/data-sharing-guidelines.

84. Schoch, C. L. et al. NCBI Taxonomy: a comprehensive update on curation, resources and tools. Database 2020, (2020).

85. NCI Thesaurus. https://ncithesaurus.nci.nih.gov/ncitbrowser/.

86. Ison, J. et al. EDAM: an ontology of bioinformatics operations, types of data and identifiers, topics and formats. Bioinformatics 29, 1325–1332 (2013).

